# Trait Heritability in Major Transitions

**DOI:** 10.1101/041830

**Authors:** M. D. Herron, S. A. Zamani-Dahaj, W. C. Ratcliff

**Affiliations:** School of Biology, Georgia Institute of Technology. North Avenue, Atlanta, GA 30332; School of Physics, Georgia Institute of Technology. North Avenue, Atlanta, GA 30332

**Keywords:** Evolution, Heritability, Major Transitions, Multicellularity, Quantitative genetics, Simulations

## Abstract

A crucial component of major transitions theory is that after the transition, adaptation occurs primarily at the level of the new, higher-level unit. For collective-level adaptations to occur, though, collective-level traits must be heritable. Since collective-level trait values are functions of lower-level trait values, collective-level heritability is related to particle-level heritability. However, the nature of this relationship has rarely been explored in the context of major transitions. We examine relationships between particle-level heritability and collective-level heritability for several functions that express collective-level trait values in terms of particle-level trait values. When a collective-level trait value is a linear function of particle-level trait values and collective size is fixed, the heritability of a collective-level trait is never less than that of the corresponding particle-level trait and is higher under most conditions. For more complicated functions, collective-level heritability is higher under most conditions, but can be lower when the environment experienced by collectives is heterogeneous. Within-genotype variation in collective size reduces collective-level heritability, but it can still exceed particle-level heritability when phenotypic variance among particles within collectives is large. These results hold for a diverse sample of biologically relevant traits. Rather than being an impediment to major transitions, we show that collective-level heritability superior to that of the lower-level units can often arise ‘for free’, simply as a byproduct of collective formation.

## Introduction

Major transitions, or evolutionary transitions in individuality, are a framework for understanding the origins of life’s hierarchy and of biological complexity [1,2]. During such a transition, a new unit of evolution emerges from interactions among previously existing units. This new unit, or collective, has traits not present before the transition and distinct from those of the units that comprise it (particles; see [3] for an in-depth discussion of collective-level traits). These collective-level traits are potentially subject to selection. Over the course of the transition, the primary level of selection shifts from the particle (lower-level unit) to the collective (higher-level unit), for example from cells to multicellular organisms or from individual insects to eusocial societies.

Evolution by natural selection requires heritable variation in phenotypes that affect fitness at the level at which selection occurs [4,5]. The breeder’s equation of quantitative genetics shows that heritability and strength of selection contribute equally to the adaptive response (see Analytical model below). When a collective-level trait is exposed to selection, it is collective-level heritability (the heritability of the collective-level trait) that determines the magnitude of the response. Collective-level heritability of traits is thus necessary for collective-level adaptations, but the emergence of collective-level heritability during a major transition has often been assumed to be difficult. For example, Michod considers the emergence of collective-level heritability through conflict mediation a crucial step in major transitions [2,6,7]. Simpson says that “From the view of some standard theory, these transitions are impossible,” in part because particle-level heritability greatly exceeds collective-level heritability [8].

Major transitions can be conceptualized as a shift from MLS1 to MLS2, in the sense of Damuth and Heisler [5], as in Okasha [9] (see also Godfrey-Smith [10], Shelton & Michod [11]). In MLS1, properties of the particles are under selection; in MLS2, it is the properties of the collectives. We follow Okasha [9] in referring to the lower-level units in a transition as ‘particles’ and the higher-level units as ‘collectives.’ Although our biological analogies are presented in terms of cells as particles and multicellular organisms as collectives, in principle our model could be extended to any pair of adjacent levels.

According to Michod [6], “…the challenge of ETI [evolutionary transitions in individuality] theory is to explain how fitness at the group level in the sense of MLS2 emerges out of fitness at the group level in the sense of MLS1.” But fitness, or selection, is only half of the breeder’s equation. Predicting the response to selection requires an estimate of heritability.

Whether or not collective-level fitness in MLS2 is a function of particle-level fitness is a matter of some disagreement (for example, Rainey and Kerr say no [11]). However, collective-level <u>phenotypes</u> must be functions of particle-level trait phenotypes, unless we accept strong emergence, a philosophical position tantamount to mysticism [13]. The function may be complex and involve cell-cell communication, feedbacks, environmental influences, etc., but it is still a function that is, in principle, predictable from particle-level trait values.

Nevertheless, the relationship between the heritability of particle-level traits and that of collective-level traits has rarely been considered in the context of major transitions, leading Okasha [14] to wonder, “Does variance at the particle level necessarily give rise to variance at the collective level? Does the heritability of a collective character depend somehow on the heritability of particle characters? The literature on multi-level selection has rarely tackled these questions explicitly, but they are crucial.” Similarly, Goodnight [15] says, “…we really do not have a good understanding of what contributes to group heritability, how to measure it, or even how to define it.”

While the role of selection has often been considered in the context of major transitions, the role of trait heritability has been relatively neglected. We examine relationships between particle-level heritability and collective-level heritability for several functions that express collective-level trait values in terms of particle-level trait values. For the simplest (linear) function, we derive an analytical solution for the relationship. For more complex functions, we employ a simulation model to explore the relationship over a range of conditions.

### Analytical model

There are several ways to estimate heritability, the proportion of phenotypic variation explained by genetic variation. If the strength of selection is known, heritability can be estimated by back-calculating from the breeder’s equation: *R* = *h*^2^*S*, where *R* is the response to selection, *S* the selection differential, and *h*^2^ the narrow-sense heritability (i.e. the proportion of phenotypic variation explained by additive genetic variation). This can be rearranged as *h*^2^ = *S/R*. Another method is to compare parent and offspring trait values: the slope of the parent-offspring regression is an estimator of heritability [16]. We use the latter method in the simulations described in the next section.

Since heritability can be defined as the proportion of phenotypic variance explained by genetic variance, one method of estimation is to partition total variance into its components using an analysis of variance. We employ this approach in an analytical model to derive the relationship between the heritability of a collective-level trait and that of the particle-level trait from which it arises. For the sake of tractability, we begin with the simplest case, assuming that the size (number of particles) of collectives is fixed and that the collective-level trait value is a linear function of the particle-level trait values. We further assume that reproduction is asexual, so the proper measure of heritability is broad-sense heritability, *H*^2^ [17]. Broad-sense heritability describes the proportion of phenotypic variation explained by all genetic variation, including both additive and non-additive components.

We imagine a population in which collectives are made up of particles and genetically distinct clones are made up of collectives. As a concrete example, we can think of a population of undifferentiated volvocine algae, such as *Gonium*, in which case the particles are cells and the collectives are colonies. Because of asexual reproduction, many genetically-identical collectives may comprise a clone. Genetic variation among clones may arise through mutation or because the population is facultatively sexual, in which case these results will only hold for evolution within the asexual phase (in the *Gonium* example, during the summer bloom that precedes autumn mating and winter dormancy).

Broad-sense heritability is the ratio of genetic variance (*V*_*G*_) to total phenotypic variance (*V*_*P*_), estimated as the ratio of among-clone variance to total phenotypic variance [17]. Inherent in this concept is that genetically identical individuals are not always phenotypically identical; *V*_*P*_ includes both genetic and non-genetic variation. Non-genetic variation can arise from maternal effects, environmental (including microenvironmental) effects, and random developmental noise. Phenotypic variation among genetically identical individuals has been extensively documented, including in bacteria [18,19], unicellular eukaryotes [20], plants [21], animals [17], and volvocine algae [22].

In this section, we use an ANOVA framework to estimate heritability as a ratio of sums of squares. Strictly speaking, heritability is a ratio of variances, not of sums of squares. However, the ratios of the relevant sums of squares converges to that of the variances as the number of categories increases (see Supplemental Information), and for all but tiny or genetically uniform biological populations, the difference between the two ratios is negligible.

Treating particles and collectives separately, the phenotype of particle *k* in collective *j* within clone *i* can be expressed as

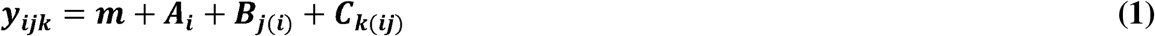

where *m* is the mean genotypic value of all clones, *A*_*i*_ is the deviation of clone *i* from *m, B*_*j*(*i*)_ is the deviation of collective *j* from the mean of clone *i*, and *C*_*k*(*ij*)_ is the deviation of particle *k* from the mean of collective *j* within clone *i*. The model in (1) describes a nested ANOVA framework, in which the sums of squared deviations from the population mean is partitioned into among-clone, among collectives within clone, and within-collective components. The among-clone component, the sum of squared deviations of *A* from *m*, is

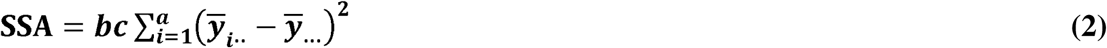

where *a, b,* and *c* are the number of clones, collectives within a clone, and particles within a collective, respectively. The sum of squared deviations of collectives within clones is

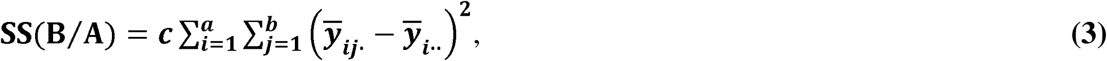

that among particles within collectives is

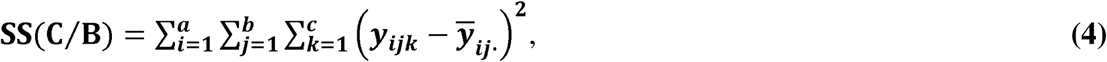

and total sum of squares is

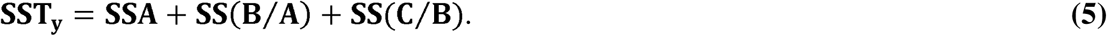

Broad-sense heritability of a particle-level trait, 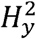, is the ratio of genetic variance to total phenotypic variance:

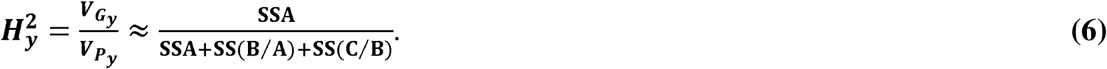

We now turn our attention to collective-level traits. The phenotype of collective *j* within clone *i* can be expressed as

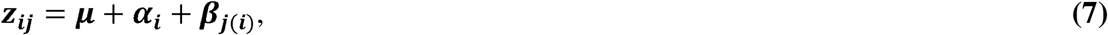

where *μ* is the mean genetic value of all clones, *α*_*i*_ is the deviation of clone *i* from *μ*, and *β*_*j*(*i*)_ is the deviation of collective *j* from the mean of clone *i*. The sum of squared deviations of *α* from *μ* is

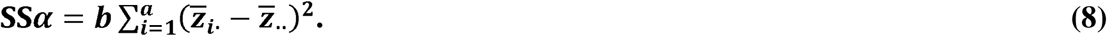

The sum of squares among colonies within clones is

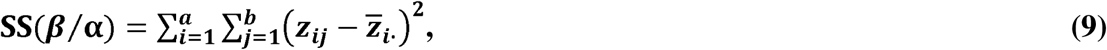

and the total sum of squares is

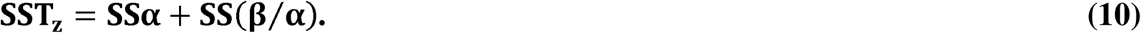

Broad-sense heritability of a collective-level trait, 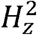, is the ratio of genetic variance to total phenotypic variance,

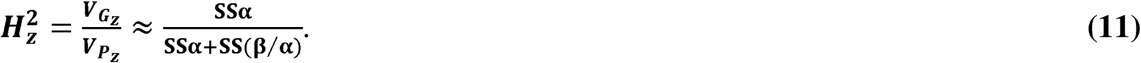

If collective-level trait value is the average of cell-level trait values, *z_ij_* = *y_ij_*·, 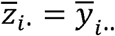, and 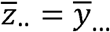. Thus SSα = *c*SSA, and SS(β/α) = *c*SS(B/A). Substituting into (11),
we get

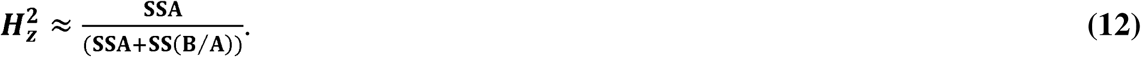

The ratio of collective-level heritability to particle-level heritability is thus

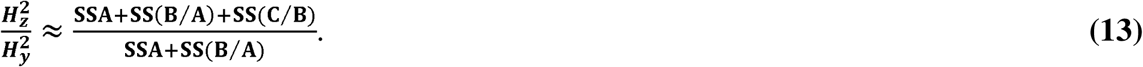

Collective-level heritability is therefore never less than particle-level heritability (i.e., the ratio of heritabilities is never less than 1), and is greater unless SS(C/B) = 0, in other words unless particles within each collective have identical phenotype.

Although we have derived this relationship assuming that the collective-level trait value is the average of particle-level trait values, the result holds for any linear function. The substitution that gets us from (11) to (12) introduces the constant *c*, which scales both numerator and denominator and therefore cancels out. Different linear functions would change the magnitude of the constant relating SSα to *c*SSA and SS(β/α) to *c*SS(B/A) but not the fact that numerator and denominator are scaled by the same constant.

The approximations in (6) and (11), which express ratios of variances as ratios of sums of squares, hold when the number of clones (*a*) and the number of genetically identical collectives within a clone (*b*) are large (Electronic Supplement 1). For example, at *a* = *b* = 10, the approximation differs from the true value by less than 1%. Thus the results of the analytical model hold for all but tiny and/or extremely genetically depauperate populations. The number of particles within a collective (*c*) does not play a role, so our results are relevant even early in a major transition, when the collectives are likely to be small. For most real biological populations, the difference between the true heritability and the sums of squares approximation will be negligible (see Electronic Supplement 1 for a simple numerical example).

### Simulation model

The correspondence between particle-level and collective-level trait values is likely to be more complicated than a linear relationship for many interesting and biologically relevant cases. Here we explore more complicated trait mapping functions using a simulation model. As above, particles grow in clonal collectives, which reproduce by forming two new collectives, each with as many particles as its parent. The initial population is founded by ten genetically distinct clones, each of which has a different genetically determined mean particle phenotype (spaced evenly between 1 and 2). These are grown for at least 7 generations, resulting in at least 127 collective-level reproductive events per genotype and 127*n* (where *n* is particle number per collective) particle-level reproductive events per genotype. Simulation models are provided as Electronic Supplements 2-8.

In this model, we consider two sources of non-genetic effects on particle phenotype (Figure 1), each of which should lower the heritability of both particle- and collective-level traits. The first is intrinsic reproductive stochasticity in particle phenotype, analogous to developmental instability [23]. In the model, we determine the phenotype of daughter cells by sampling from a distribution centered on the parent’s genetic mean, with standard deviation σ. As shown in the analytical model above, by averaging out this variation, collectives can gain a heritability advantage over cells.

**Figure 1.**
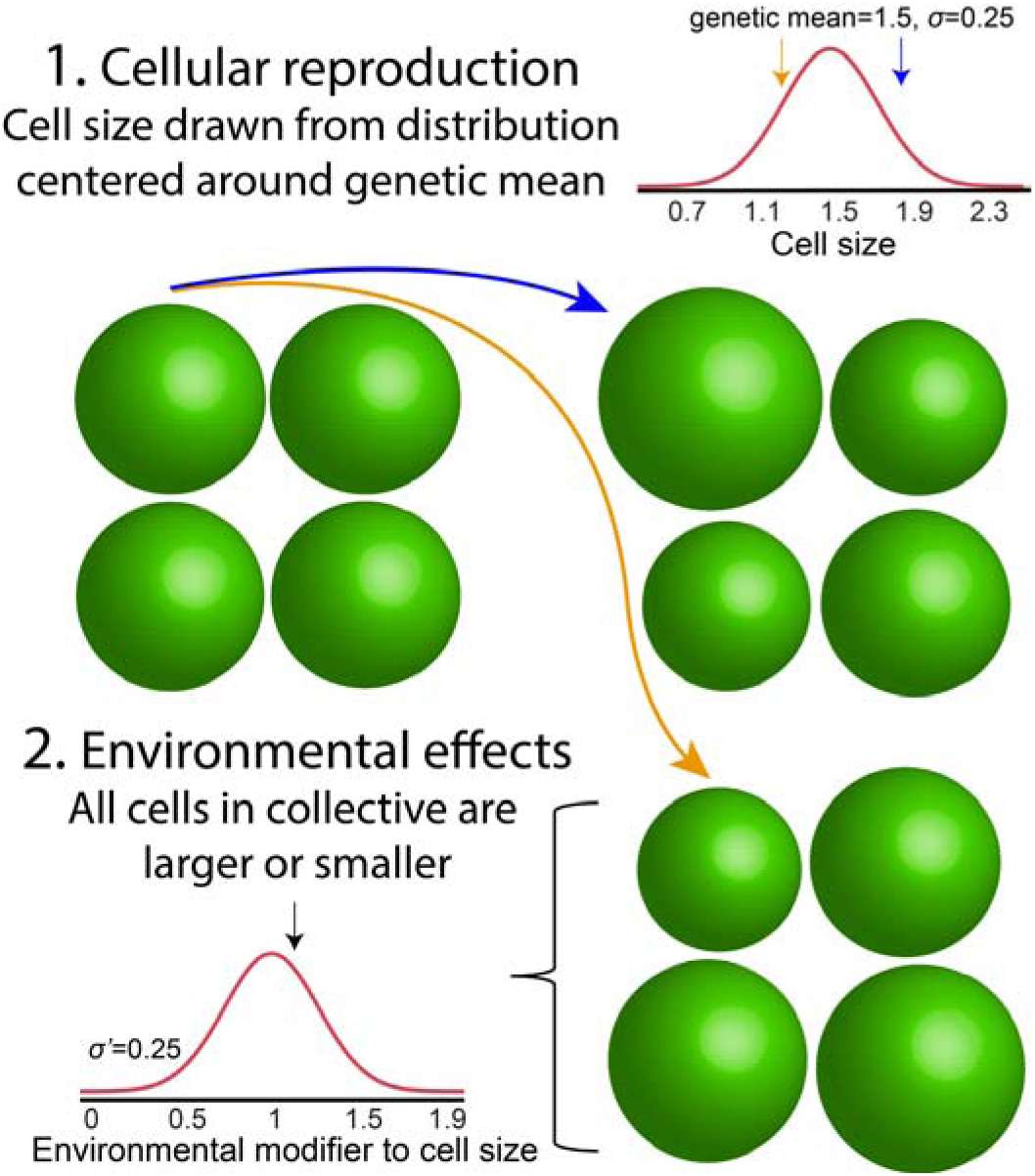
Two non-genetic modifiers to cell phenotype. There are two non-genetic influences on particle phenotype (cell size in this example) in our model: developmental instability, a stochastic effect that varies a particle’s phenotype from its genetic mean (with standard deviation σ), and environmental effects, which modify the phenotype of all particles in a collective by the same amount (with standard deviation σ□).

Our simulation also considers the phenotypic effects of environmental heterogeneity. Here, we model collectives as independently experiencing different environmental conditions that affect the phenotypes of all cells within them in the same manner. To extend the biological analogy offered above, *Gonium* colonies growing near the surface of a pond (where light and CO_2_ are abundant) may form colonies with larger cells than clonemates near the bottom. We implemented this in our model by assigning a size modifier, drawn from a normal distribution centered on 1 with standard deviation σ□, to each collective. We then multiplied the phenotype of each particle within the collective by this modifier. This source of phenotypic heterogeneity should reduce the heritability of collectives more than particles, simply because collectives experience a relatively higher frequency of stochastic events than particles do (each collective gets assigned a different size multiplier, but every particle within that collective experiences the same size multiplier).

We examine the effect of each of the above sources of phenotypic variation independently for the example of cells (particles) within nascent multicellular organisms (collectives). For a linear relationship, collective size is simply the sum of the sizes of cells within the collective. For both cells and collectives, heritability is assessed by calculating the slope of a linear regression on parent and offspring phenotype [16]. In this simple case, mean collective-level heritability is always greater than or equal to cell-level heritability. Only when σ = 0 (*i.e.*, when all cells within a collective have identical phenotype) are cell- and collective-level heritability equal, in agreement with the analytical model. Greater developmental instability for cell size increases the advantage of collective-level heritability over cell-level heritability (Figure 2a). Larger collectives, which average out cellular stochasticity more effectively, experience a greater increase in heritability than smaller collectives (Figure 2a). Note that the simulations run in Figure 2a reflect a very patchy environment in which environmental effects on cell size within collectives are large (σ□ *=* 0.25). While our model is not explicitly spatial, when σ□ is high, different collectives experience different environmental effects on their mean cell size, simulating the effects of a patchy environment. Increasing the magnitude of these environmental effects on cell size diminishes the difference in heritability between collectives and cells, but mean collective-level heritability is still greater than cell-level heritability for all parameter combinations (Figure 2b).

**Figure 2.**
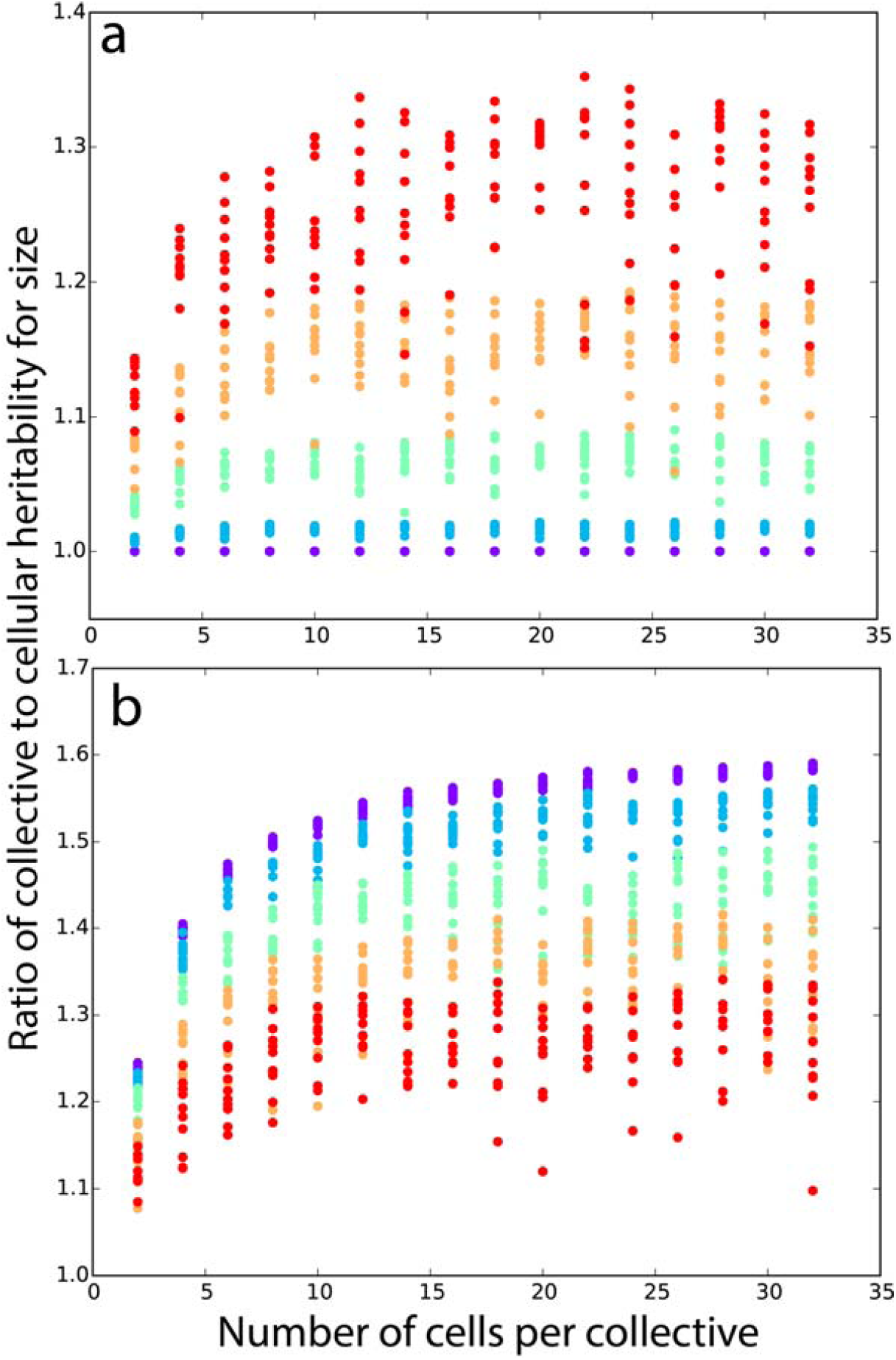
Collective-level heritability of size is greater than particle-level heritability for size. In **a**), we hold the effect of the environment fixed (standard deviation σ□ = 0.25), and vary the degree of developmental instability for particle size σ: 10^-4^ (purple), 0.0625 (blue), 0.125 (green), 0.1875 (yellow), 0.25 (red). In the absence of developmental instability for size, collective and cell-level heritabilities are identical. Greater developmental instability increases relative collective-level heritability. **b**) Here we hold developmental instability fixed at σ = 0.25, and vary between-collective environmental effects on cell size from σ□ = 10^-4^ (purple) to 0.25 (red). When developmental instability is nonzero, larger collectives improve collective-level heritability. We ran ten replicates of each parameter combination and simulated populations for nine generations of growth.

The volume of the cellular collective (Figure 2, Figure 3a), which is simply the sum of the cell volumes within it, represents the simplest function mapping cellular to multicellular trait values. We now consider more complicated nonlinear functions relating cellular to multicellular trait values, some of which have biological relevance to the evolution of multicellularity. For each function, we calculated the relative heritability of collective-to cell-level traits for 32-celled collectives across 1024 combinations of σ and σ□ ranging from 0 to 0.25.

**Figure 3.**
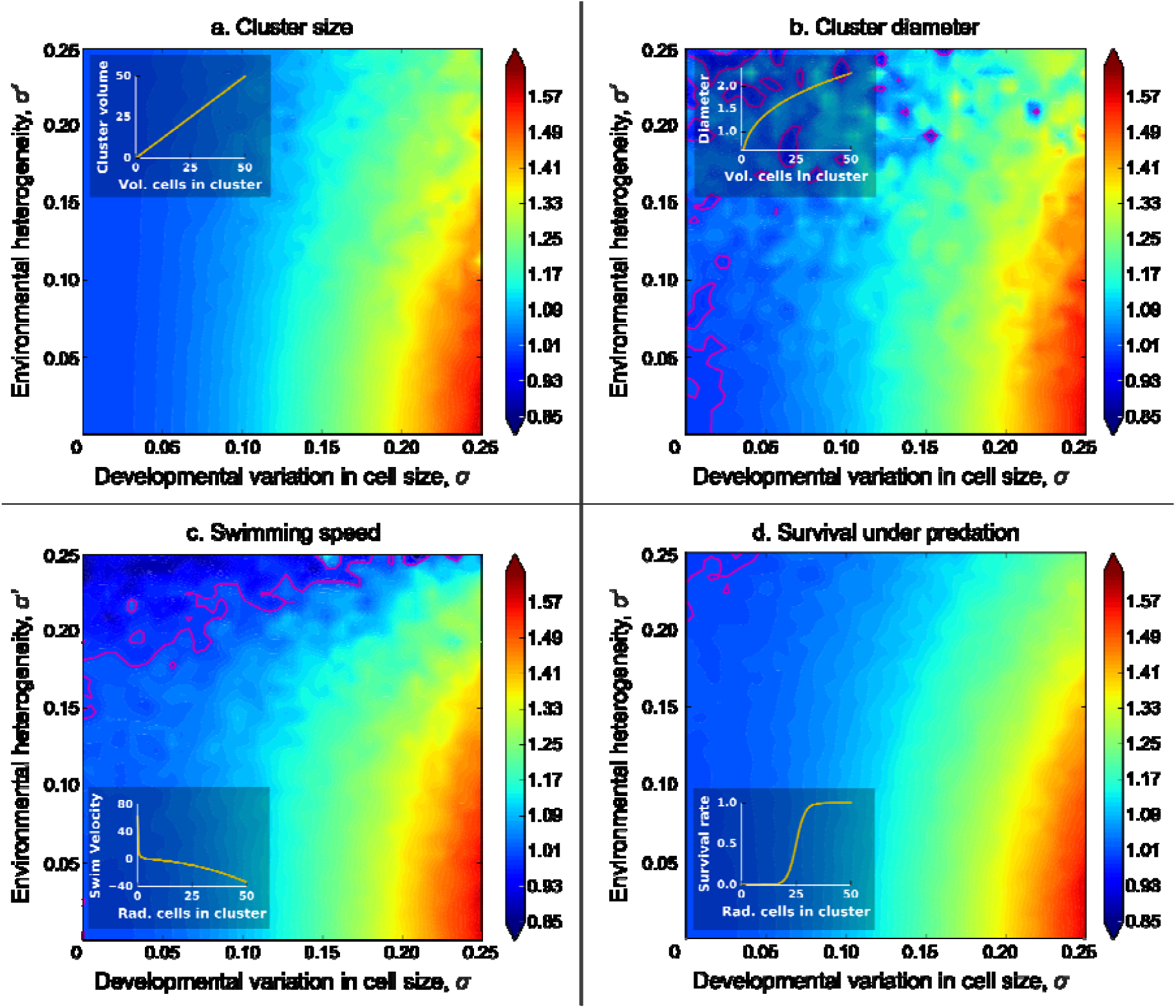
Relative heritability of various collective-level traits to cell-level heritability for size. Here we examine the heritability of four multicellular traits that depend on the size of their constituent cells, relative to cellular heritability for size. The relationship between the size of the cells within collectives and the multicellular trait are shown as insets. We consider three biologically-significant traits with different functions mapping the size of cells within the collective onto collective phenotype. The heritability of collective size (**a**) and diameter (**b**) is always higher than cell-level heritability for size, and is maximized when cellular developmental noise is greatest and among-collective environmental effects are smallest (lower right corner). We modeled swimming speed (**c**) based on the model of Solari *et al.* (2006) for volvocine green algae. We also considered survival rate under predation as a logistic function of radius (**d**). Like a and b, collective-level heritability is highest relative to cell-level heritability when environmental heterogeneity is minimal. Pink contours denote relative heritability of 1. In these simulations we consider 32 cell collectives grown for 7 generations. The colormap denotes collective-level heritability divided by cell-level heritability for size across 1024 σ, σ□ combinations.

The first nonlinear collective-level trait we consider is its diameter. Large size is thought to provide a key benefit to nascent multicellular collectives when they become too big to be consumed by gape-limited predators [24,25]. For a collective that is approximately spherical, the trait that actually determines the likelihood of being eaten is diameter, which is therefore an important component of fitness. For geometric simplicity we assume that the cells within the collective are pressed tightly together into a sphere, allowing us to calculate collective radius as 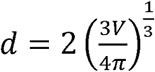, where *V* is the sum of the cell volumes within the collective. Collective volume (Figure 3a) and diameter (Figure 3b) exhibit similar dynamics, with collective-level heritability always exceeding cell-level heritability, and being maximized under conditions of strong cell size stochasticity (high σ) and no environmental heterogeneity (low).

Next, we consider swimming speed as a function of cell radius. We based this simulation on the hydrodynamics model of volvocine green algae derived by Solari *et al.* [26]. For simplicity, we modeled 32-celled, undifferentiated collectives (GS colonies in [26]), which would be similar to extant algae in the genus *Eudorina*. Given these assumptions, the function relating cell radius to upward swimming speed (Equation 4 from [26]) can be simplified to

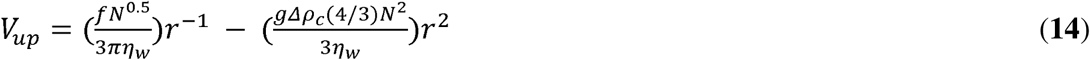

where *f* is average effective upward swimming force per cell, *N* is the number of cells per collective, *ηw* is water viscosity, *r* is the average radius of cells in the collective, and *Δρc* is the density difference between cells and water. Electronic Supplement 9 provides a more detailed description of the derivation of Equation 14.

Using the numerical values in Solari *et al* [26], *ηw* = 0.01 g/cm·s, *Δρc* = 0.047g/cm^3^, and *f* = 2.4 × 10^-7^ g·cm/s^2^, so

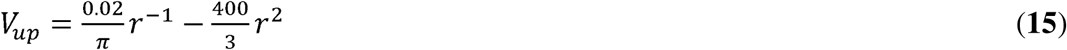

In this model, the swimming force of cells is independent of cell size, so, as cells get larger the collective will become heavier (more negatively buoyant) without a corresponding increase in total swimming force, and therefore its upward swimming speed will decrease. Thus upward swimming speed is a monotonically declining function of cell radius (Fig. 3c inset), unlike the functions for volume and diameter (Fig. 3a, 3b insets), both of which are monotonically increasing. Nevertheless, the general behavior of heritability is very similar to the previous ones and for a wide range of parameter values, the collective-level trait has a higher heritability than the cell-level trait (Fig. 3c).

Next, we consider a function describing a collective’s survival rate in the presence of a predator that can only consume collectives below a certain size. We calculated the survival rate (*c*) as a logistic function of the collective’s radius, effectively assuming that predation efficiency drops off quickly when collectives reach a threshold size (Fig. 3d inset):

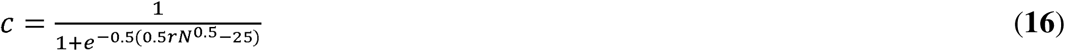

As with the previous functions (Fig. 3a-c), collective-level heritability is greater than cell-level heritability for much of the trait space and is maximized under conditions of high cellular stochasticity (σ) and low environmental heterogeneity (σ□; Fig. 3d).

Finally, we consider the case in which the simplifying assumption of constant cell number does not hold. Instead, the number of cells per collective fluctuates around the genetic mean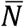. In this case, each collective reproduces two new collectives, but the number of cells per new collective is a random variable drawn from a normal distribution with mean 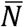 and coefficient of variation *CV*_*N*_ (the coefficient of variation for a normal distribution is the ratio of standard deviation to the mean). We chose to represent variation in the number of cells per collective as *CV*_*N*_ instead of standard deviation so that the range of variation would not change with the size of the collective.

Variation in cell number, unlike the developmental and the environmental variation, does not affect the heritability of cells, only that of collectives. Therefore, we expected that increasing *CV*_*N*_ would decrease the ratio of collective-level to cell-level heritability. To test this effect, we calculated the relative heritability of size (volume) for collectives and cells across 1024 combinations of and *CV*_*N*_ ranging from 0 to 0.25 (). The simulation shows that the *CV*_*N*_ has a strong effect on collective-level heritability (Fig. 4). As *CV*_*N*_ increases, the ratio of collective-to cell-level heritabilities decreases, falling below one when the magnitude of is similar to or smaller than that of *CV*_*N*_ (Figure 4).

**Figure 4.**
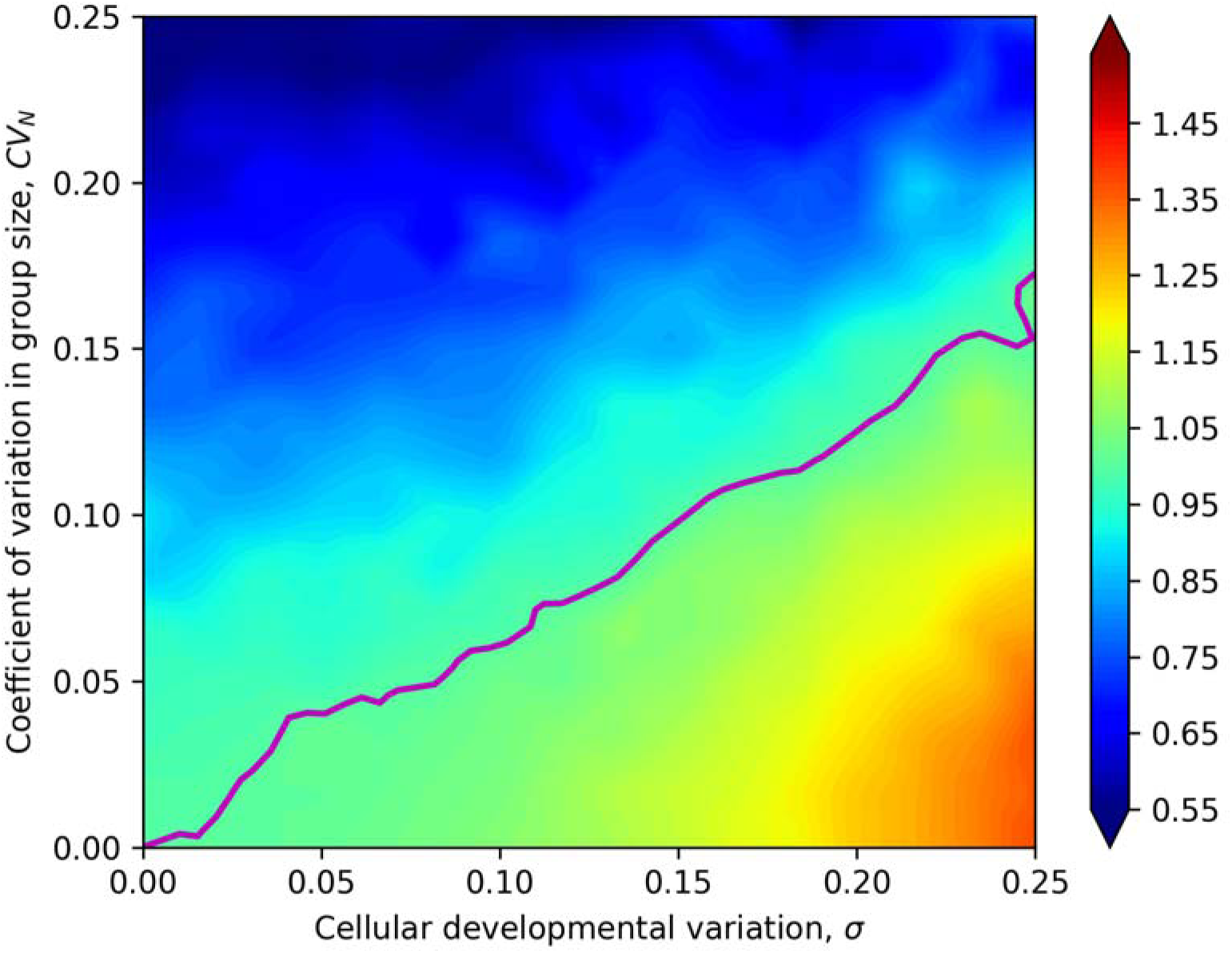
Relative heritability of collective size to cell size when the number of cells per collective varies. When the coefficient of variation for cell number per collective (CV_N_) is low, collective-level heritability is always higher than cell-level heritability, but this advantage is undercut by increased variation in cell number. The ratio of collective-to cell-level heritability is maximized when developmental variation in cell size (σ) is large and variation in the number of cells per collective is low. The pink contour denotes a ratio of collective-level to cell-level heritability of 1. In these simulations, we consider collectives with a genetic mean of 32 cells grown for 7 generations. The colormap denotes collective-level heritability divided by cell-level heritability for size across 1024 σ, *CV*_*N*_ combinations.

## Discussion

Using a quantitative genetics framework, we have derived an analytical solution for the relationship between particle-level and collective-level heritability for a limited case. When particle number is constant and the collective-level trait value is a linear function of the particle-level trait values, the organismal heritability turns out to be a simple function of the cell-level heritability. In contrast to claims that particle-level heritability is always higher than collective-level heritability (e.g. [8]), we have shown that collective-level heritability is higher over a wide range of conditions. Because this result depends on the number of clones and the number of colonies within a clone, it may not hold for very small populations or those with little genetic variation. This is not a major limitation, though, since tiny, genetically homogeneous populations are unlikely to be the ones experiencing selectively driven evolutionary transitions in individuality.

This analytical result is a step toward understanding the relationship between heritabilities at two adjacent hierarchical levels, but the assumptions of constant particle number and linear function are restrictive. The simulation model shows that the results are somewhat dependent on the function relating the trait values at the two levels. However, these functions were chosen to be diverse, and the behavior of the relative heritabilities is nevertheless qualitatively similar, increasing with cellular developmental variation (σ), decreasing with environmental heterogeneity (σ□), and exceeding 1 for most of the parameter space.

Of course, we have not (and cannot) comprehensively explored the universe of possible functions relating collective-level traits to particle-level traits. What we have done is explore a small sample of this space, with functions ranging from extremely simple (volume) to somewhat more complex (swimming speed, survival under predation). We do not claim that the high heritabilities estimated for these collective-level traits would apply to all such traits, and a full accounting of possible functions is beyond the scope of this (or any) study. Rather, we have shown that for at least some such functions, the resulting collective-level traits can have high heritability, and thus be altered by selection, early in an evolutionary transition in individuality.

All four of the collective-level traits in the simulation models are potentially biologically relevant. Volume and diameter are both aspects of size, which can be an important component of fitness both in evolutionary transitions in individuality [27] and in life history evolution [28]. Swimming speed is a measure of motility, which has selective consequences for a wide range of organisms, including many animals and microbes. For planktonic organisms, a positive upward swimming speed provides active control of depth, allowing some control over light intensity (for autotrophs) and prey abundance (for heterotrophs). Survival under predation obviously has important fitness implications for many organisms, and both theoretical and experimental evidence implicate predation as a possible selective pressure driving the evolution of multicellularity. Kirk, for example, suggests that a “predation threshold” above which algae are safe from many filter feeders may have driven the evolution of multicellularity in the volvocine algae [29]. Microbial evolution experiments in the algae *Chlorella* and *Chlamydomonas* have shown that predation can drive the evolution of undifferentiated multicellular clusters [30–32].

In our simulations, we examined the effects of three independent sources of phenotypic variation affecting the relative heritability of particle and collective-level traits. Stochastic variation in cell size around the clone’s genetic mean (σ) reduces the absolute heritability of cells and collectives by introducing non-heritable phenotypic variation. By averaging across multiple cells, however, collectives reduce the effects of this phenotypic variation, providing them with a relative heritability advantage over cells.

We also considered the effect of environmental heterogeneity in which all of the cells within a collective are affected in the same manner (σ*’*). Collectives are disproportionately affected: each collective is assessed a different size modifier, but all of the cells within these collectives are affected in the same manner. As a result, collectives experience *n-*fold more stochastic events (where *n* is the number of cells per collective), which reduces their heritability relative to cells. The influence of these sources of variation is evident in the contour plots of Figure 3: the relative heritability of collectives to cells is maximized when cellular stochastic variation is high and environmental heterogeneity low (lower right corner of the plots). The effect of environmental heterogeneity in our simulations is consistent with the empirical finding of Goodnight [33] that group selection of *Arabidopsis* was more effective when among-deme environmental variance was low.

Finally, we considered variation in the number of particles per collective. Such variation substantially reduces the heritability of a collective-level trait. Even with reasonably large variation in collective size, though, the collective-level trait retains most of the heritability of the particle-level trait on which it is based (for example, ∼55% at a CV_N_ in particle number of 0.25).

Our results differ from previous considerations of heritability in important respects. For example, Queller [34] presents a useful reformulation of the Price equation for selection at two levels:

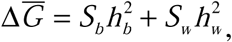

in which 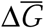 is the change in average trait value, *S*_*b*_ and *S*_*w*_ are the selection differentials between collectives and within collectives, respectively, and *h*^2^_*b*_ and *h*^2^_*w*_ are the heritabilities of the collective-level and individual-level traits, respectively. This formulation partitions the response to selection on a particle-level trait into within- and among-collective change, but the focus is still on particle-level traits. Our focus is on the evolution of collective-level traits. In the terminology of Damuth and Heisler [5], our focus is on MLS2, while Queller’s is on MLS1. In addition, Queller makes no attempt to derive the relationship between collective-level heritability and particle-level heritability.

Michod and Roze [2] have previously modeled the relationship between particle-level and collective-level heritability of fitness during a major transition. However, as Okasha [14] points out, heritability of fitness only ensures that mean population fitness will increase over time. For selection to result in directional phenotypic change, it is phenotypes that must be heritable. Futhermore, Michod and Roze focused on within-organism genetic change. Our models assume that such change is negligible, as is likely to be true early in a transition, when collectives (*e.g.*, nascent multicellular organisms) presumably include a small number of clonally-replicating particles (*e.g.*, cells).

Okasha [35] considers heritability in MLS1 (which he refers to as group selection 2) and MLS2 (his group selection 1) but does not attempt to derive a relationship between heritabilities at two levels. We have focused on just this relationship, because knowing the ratio of heritabilities is necessary to predict the outcome of opposing selection at two levels. This has important implications for collective-level traits that arise from cooperation among particles. The presumed higher heritability of the particle-level traits has been seen as a problem for the evolution of cooperation that benefits the collective [2,8,36–38]. Our results show that this problem does not always exist.

Several previous papers have shown that group-level heritability (collective-level heritability in our terminology) exists and can be substantial. Slatkin [39], for example, showed that one measure of group-level heritability, fraction of total variance between lines, is substantial both in an additive model and in the *Tribolium* experiments of Wade and McCauley [40]. Under some conditions, the between-line variance of a linear trait such as the one we consider in our analytical model exceeds the within-line variance.

Bijma, Wade and colleagues [41–43] showed that variance in the total breeding value of a population can be increased, even to the point of exceeding phenotypic variance, by interactions among individuals. Our model does not consider (or require) interactions among individuals. Further, their model and empirical example are exclusively concerned with individual-level traits (particle-level traits in our terminology), for example survival days in chickens. They do not estimate group heritability as such, and judge that “it is unclear how this parameter should be defined or estimated.”

Goodnight [15] considers the ratio of group-level heritability to individual-level heritability (in the narrow sense) using contextual analysis. Although this paper does not provide a formula to calculate this ratio, its inequality sets a minimum bound (with the assumption that selection at the two levels is in opposition). As in our analyses, Goodnight shows that group-level heritability can exceed individual-level heritability in some circumstances.

Several simplifying assumptions underlie our models, most importantly the genetic identity of particles within collectives. This condition only applies to a subset of the major transitions. Queller recognized two subcategories within Maynard Smith and Szathmáry’s [1] list of transtions, which he called “egalitarian” and “fraternal” transitions [44]. Briefly, egalitarian transitions involve particles that may be genetically distinct, or even from different species, such as the alliance of a bacterium with an Archaean that gave rise to the eukaryotic cell. Fraternal transitions are those in which the particles are genetically similar or identical, such as the origins of eusociality and of most multicellular lineages.

Because of our assumption of genetic identity among particles, we cannot generalize our results to all types of major transitions. Egalitarian transitions will not normally meet this criterion. A possible exception is aggregative multicellularity, as seen in cellular slime molds and myxobacteria, when assortment is so high that fruiting bodies are genetically uniform. This is probably uncommon [45], but it does happen [46,47]. Transitions in which reproduction of particles is obligately sexual, such as the origins of eusociality, also violate this assumption.

A better fit for our models is clonal multicellularity, which is probably the most common type of major transition. An incomplete list of independent origins of clonal multicellularity includes animals; streptophytes; chytrid, ascomycete, and basidiomycete fungi; florideophyte and bangiophyte red algae; brown algae; peritrich ciliates; ulvophyte green algae; several clades of chlorophyte green algae; and filamentous cyanobacteria [48–51]. In most cases the early stages in these transitions probably violated the assumption of uniform particle number per collective, but our simulations show that our main results are robust to reasonable violations of this assumption.

One example that does approximate all of our assumptions is that of the volvocine green algae, an important model system for understanding the evolution of multicellularity. Volvocine algae undergo clonal reproduction only occasionally punctuated by sex, are small enough that within-collective mutation probably has negligible phenotypic effects, and have cell numbers that are under tight genetic control.

## Conclusion

A great deal of work has gone into understanding the selective pressures that may have driven major evolutionary transitions. However, heritability is just as important as the strength of selection in predicting evolutionary outcomes. We have shown that, given some simplifying assumptions, heritability of collective-level traits comes ‘for free’; that is, it emerges as an inevitable consequence of group formation. Qualitatively, this result holds across a wide range of parameters and for a diverse sample of biologically relevant traits. Collective-level heritability is maximized (relative to particle-level heritability) when phenotypic variation among particles is high and when environmental heterogeneity and variation in collective size are low. Understanding the emergence of trait heritability at higher levels is necessary to model any process involving multilevel selection, so our results are relevant to a variety of other problems.

## Declarations

### Ethics approval and consent to participate

Not applicable

### Consent for publication

Not applicable

### Availability of data and material

All data generated or analysed during this study are included in this published article [and its supplementary information files].

### Authors’ contributions

MDH conceived the project, developed the analytical model, contributed to the simulation models, and contributed to writing the manuscript. SAZ-D and WCR developed the simulation models and contributed to writing the manuscript. All authors read and approved the final manuscript.

## Acknowledgements

We would like to thank Sam Brown, Peter Conlin, Michael Doebeli, Rick Michod, and Deborah Shelton for helpful discussions.

